# The extracellular ATP receptor P2RX7 imprints a pro-memory transcriptional signature in effector CD8^+^ T cells

**DOI:** 10.1101/2021.05.25.445679

**Authors:** Trupti Vardam-Kaur, Sarah van Dijk, Changwei Peng, Kelsey Wanhainen, Stephen C. Jameson, Henrique Borges da Silva

## Abstract

Development of central memory (Tcm) and resident memory (Trm) CD8^+^ T cells, which respectively promote immunity in the circulation and in barrier tissues, is not completely understood. Tcm and Trm cells may arise from common precursors, however their fate-inducing signals are elusive. We found that virus-specific effector CD8^+^ T cells display heterogeneous expression of the extracellular ATP sensor P2RX7. P2RX7-high expression is confined, at peak effector phase, to CD62L^+^ memory precursors which preferentially form Tcm cells. Among early effector CD8^+^ T cells, asymmetrical P2RX7 distribution correlated with distinct transcriptional signatures, with P2RX7-high cells enriched for memory and tissue-residency sets. P2RX7-high early effectors preferentially form both Tcm and Trm cells. Defective Tcm and Trm formation in P2RX7 deficiency is significantly reverted when the transcriptional repressor Zeb2 is ablated. Our study indicates that unequal P2RX7 upregulation in effector CD8^+^ T cells is a foundational element of the early Tcm/Trm fate.

## Introduction

The development of immunological memory is paramount for host protection against secondary pathogen exposure^1^. The establishment of antigen-specific cytotoxic memory CD8^+^ T cells is specifically important for antiviral protective immunity^2^. Over the past two decades, it has become clear that CD8^+^ T cell memory is comprised of distinct cell subsets with different migration, function, and longevity properties^3^. These characteristics revealed four major memory CD8^+^ T cell subsets. Long-lived effector cells (LLECs) are mostly confined to blood and blood-associated tissues^4, 5^, while effector memory cells (Tem) recirculate between secondary lymphoid organs, blood, and non-lymphoid tissues^4^. Two other subsets display heightened longevity, secondary response potential and stemness. Central memory cells (Tcm) which are mostly present in secondary lymphoid organs and express the defining marker L-selectin (CD62L)^6^, and tissue resident memory cells (Trm) which are prominent in non-lymphoid tissues and, in normal circumstances, do not recirculate^7, 8, 9^. Tcm and Trm cells, although present in distinct environments, share many transcriptional and functional characteristics^10^, and may arise from common precursors^11^. Previous studies suggest these common precursors are found at the early effector immune response, express low levels of the terminal effector (TE)-linked molecule Killer cell lectin-like receptor subfamily G member 1 (KLRG1), and high levels of the stemness-associated transcription factor T cell factor 1 (TCF-1)^12, 13, 14, 15^. While considerable advances on the transcriptional and metabolic control of early memory precursors have been made^13, 14, 15, 16, 17^, less is known about whether and how extracellular signals contribute to early diversion toward the memory fate. Understanding how this occurs will help identify factors simultaneously promoting Tcm and Trm cells.

Many extracellular signals can be sensed by antigen-specific CD8^+^ T cells, such as cytokines (e.g. IL-7, IL-2, IL-15, IFN-γ) and chemokines (e.g. CXCL9, CXCL10) that are mostly produced by antigen-presenting cells (APCs) or stromal cells^4, 18, 19, 20^. Other signals can be released passively by the inflamed tissue or infected cells, such as nucleotides (e.g. ATP)^21^. Extracellular ATP (eATP) can be sensed by CD8^+^ T cells through the low-affinity ion channel P2RX7^22^. In recent studies, we have defined that CD8-intrinsic P2RX7 is required for the establishment of both Tcm and Trm cells^23, 24^. P2RX7 promotes metabolic adaptations fundamental for memory CD8^+^ T cell differentiation, namely activation of the AMPK signaling pathway and promotion of mitochondrial function^23^. We have also found that P2RX7 promotes the upregulation of the TGF-β signaling pathway^24^, which is crucial for epithelial Trm populations^25^. Both Tcm and Trm cells express high levels of and require P2RX7, suggesting two possible scenarios on how P2RX7 simultaneously promote these subsets. First, P2RX7 stimulation may promote survival of Tcm and Trm cells after they have differentiated, sustaining each subset independently. Our previous results suggest that, at least partially, this is the case, since continued P2RX7 expression is needed for Tcm and Trm maintenance^24^. Alternatively, intracellular pathways affected by P2RX7 may be induced in early effector CD8^+^ T cells with the potential to become either Tcm or Trm – i.e., P2RX7 signaling could promote the arising of early memory precursors. Indeed, the role of P2RX7 in upregulation of the TGF-β signaling pathway occurs at the early effector phase^24^.

In this study, we report that P2RX7 simultaneously promotes establishment of Tcm and Trm cells through positive control of a subset of early memory precursors. We show that P2RX7 expression at the early effector phase is not homogeneous, and that early effector CD8^+^ T cells expressing higher levels of P2RX7 preferentially form long-lived Tcm and Trm populations. Performing a transcriptional comparison between P2RX7 high-versus low-expressing early effector CD8^+^ T cells, we find that many pro-memory and pro-residency gene signatures are enriched in P2RX7-expressing early effector CD8^+^ T cells. Finally, we provide evidence that P2RX7-mediated downregulation of the transcriptional repressor Zeb2 aids in the acquisition of an early memory precursor phenotype and subsequent Tcm and Trm cell establishment. These results indicate that the eATP receptor P2RX7 plays a crucial role in the initial differentiation of Tcm/Trm common memory precursors, serving as an important extracellular environment sensor inducing a memory fate in effector CD8^+^ T cells.

## Results

### P2RX7 expression is heterogeneous within CD8^+^ memory precursors and correlates with pro-memory molecules

In recent studies, we have found that, in response to systemic viral infections such as LCMV, P2RX7 is quickly upregulated in effector CD8^+^ T cells^24^. At peak effector phase, P2RX7 is preferentially expressed by memory precursor CD8^+^ T cells (MPs) if compared to terminal effectors (TEs)^23^ (Fig. 1a). The expression pattern of P2RX7 within MPs, however, is not homogeneous, being found instead as a continuum (Fig. 1b). An arbitrary division between MPs expressing the highest P2RX7 (P2RX7^hi^) and the lowest P2RX7 (P2RX7^lo^) revealed profound differences in their phenotypes. CD62L expression, which marks Tcm precursors^13^ in approximately 2% of antigen-specific CD8^+^ T cells and are preferentially found in MPs (Fig. S1a), is almost exclusively confined to P2RX7^hi^ MPs (Fig. 1b). In addition, the expression of many other proteins preferentially expressed by memory CD8^+^ T cells are higher in P2RX7^hi^ MPs, such as CD44, CD101, TCF-1 (Fig. 1c-d), CXCR5 (Fig. S1b) and even the MP-defining molecule CD127 (Fig. S1c). Expression of CD8α itself is also preferentially found in P2RX7^hi^ MPs, although the same pattern can be also seen when TEs are divided in P2RX7^hi^ (less than 5% of TEs – data not shown) and P2RX7^lo^ (Fig. S1d). Conversely, expression of the exhaustion/TE marker Tim3 is preferentially found in P2RX7^lo^ MPs (Fig. 1c-d). All these expression patterns are also true when adoptively transferred, LCMV-specific P14 cells are assessed (Fig. S1e). Overall, our data shows that the MP subset display significant phenotypic heterogeneity, which can be defined by P2RX7 expression level.

**Figure 1.**
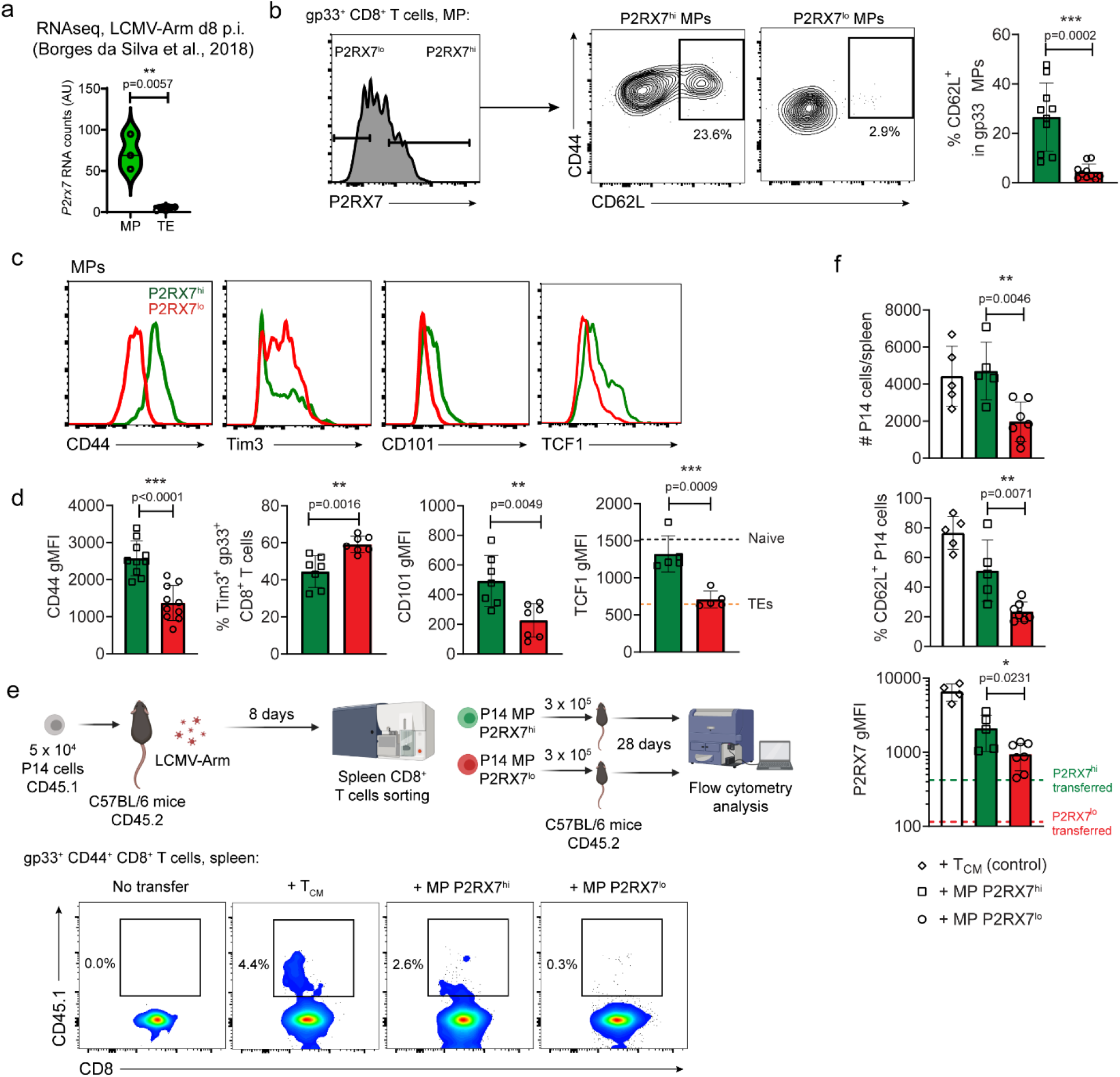
P2RX7 expression is heterogeneous among memory precursors and favor Tcm generation. **(a)** *P2rx7* mRNA counts from an RNA-seq comparison between MPs and TEs done by our group in a previous study. **(b-d)** C57BL/6 mice were infected with LCMV-Arm and harvested at 8 days post-infection; the protein expression of multiple molecules in antigen-specific (gp33-tetramer^+^) CD8^+^ MPs is shown. **(b)** Left: representative histogram showing P2RX7 expression in gp33^+^ MPs. Center: Representative contour plots showing the expression of CD44 and CD62L in P2RX7^hi^ vs P2RX7^lo^ MPs. Right: Average percentages of CD62L^+^ cells within P2RX7^hi^ (green, open squares) and P2RX7^lo^ (red, open circles) MPs. **(c)** Representative histograms of the expression of CD44, Tim3, CD101 and TCF1 in P2RX7^hi^ (green) and P2RX7^lo^ (red) MPs. **(d)** Geometric mean average values (gMFI) of CD44, CD101 and TCF-1, and percentages of Tim3^+^ in P2RX7^hi^ (green, open squares) and P2RX7^lo^ (red, open circles) MPs. **(e)** P14 cells (CD45.1^+^) were transferred to C57BL/6 (CD45.2^+^) mice which were subsequently infected with LCMV-Arm. After 8 days, P2RX7^hi^ P14 MPs (30% highest) and P2RX7^lo^ P14 MPs (30% lowest) were sorted, and adoptively transferred into infection matched C57BL/6 mice (3 × 10^5^ cells/mouse). After 28 days, the numbers and phenotype of transferred P14 cells were assessed in spleens of recipient mice using flow cytometry. On the bottom, representative plots showing the P14 populations present in recipient mice, including negative (No transfer) and positive (+ Tcm) controls. **(f)** Numbers of transferred P14 cells/spleen (top left), percentage of CD62L^+^ P14 cells (bottom left), and P2RX7 gMFI (top right) in recipient mice for positive control (Tcm) cells (white, open diamonds), P2RX7^hi^ MPs (green, open squares), and P2RX7^lo^ MPs (red, open circles). (**a, d, f**) – Unpaired t-test, * - p<0.05, ** - p<0.01 and *** - p<0.001. (**b-f**) Average values ± SD, data pooled from 3 independent experiments.

### High P2RX7 expression in MPs favor differentiation of Tcm cells

The heterogeneity found within MPs suggested that P2RX7 expression levels may correlate with differentiation into long-lived memory T cells. To assess this, we sorted P2RX7^hi^ and P2RX7^lo^ MP P14 cells, and performed secondary transfers into infection-matched mice (Fig. 1e). As positive controls we sorted and adoptively transferred central memory (Tcm) P14 cells, which efficiently maintained in recipient mice upon secondary transfer (Fig.1e-f)^26^. Within the MP pool, P2RX7^hi^ cells were present in significantly higher numbers compared to P2RX7^lo^ counterparts (Fig. 1e-f), and displayed increased proportion of CD62L-expressing cells, i.e. a Tcm phenotype (Fig. 1f). The expression of P2RX7 remained higher in P2RX7^hi^ MP progenies (Fig. 1f), but even within the fewer P2RX7^lo^ MP progenies, we found increased P2RX7 expression if compared to initial expression levels on donor cells (Fig. 1f). Overall, these data indicate that higher P2RX7 expression in MPs favor the establishment of memory CD8^+^ T cells, especially the Tcm subset.

### Qualitative and quantitative differences in P2RX7 expression correlate with a pro-memory transcriptional and protein signature in early effector CD8^+^ T cells

Our findings within the MP subset strongly suggests that heterogeneity in P2RX7 expression helps define CD8^+^ T cells that preferentially establish memory subsets. However, Trm cells do not exclusively arise from MPs, rather being derived from precursors found in the circulation at the early effector phase^27^. P2RX7 expression is also heterogeneous in spleen early effector CD8^+^ T cells (Fig. S2a), thus we hypothesized that P2RX7 levels would also correlate with the expression of pro-memory markers. First, we conducted a transcriptional assessment of whether the presence of P2RX7 in early effectors would be associated with a pro-memory signature. RNAseq analysis of WT versus *P2rx7*^-/-^ early effector P14 cells revealed >200 differentially expressed genes (DEGs) (Fig. 2a-b, Table S1). WT P14 early effectors are enriched for many genes usually found in memory CD8^+^ T cells, e.g. *Cd101, Bach2, Ccr7, Cxcr5, Tcf7*^13,28^ (Fig. 2b, Table S1). Simultaneously, genes preferentially associated with a Trm phenotype such as *Itgae*^29^ are enriched in WT early effectors (Fig. 2b, Table S1). Conversely, *P2rx7*^-/-^ early effectors are enriched for either TE-linked genes (e.g. *Cx3cr1, Klrg1, Zeb2*)^13,28^ or genes typically downregulated in Trm cells such as *S1pr1* and *S1pr5*^29, 30^ (Fig. 2b, Table S1). Predictably, many pathways associated with memory CD8^+^ T cells such as stemness-related are enriched in WT early effectors (Fig. S2b-c). Simultaneously, apoptosis-related pathways and pro-migration (i.e. anti-residency) pathways are enriched in *P2rx7*^-/-^ early effectors (Fig. S2b-c, Table S2). Accompanying these characteristics, GSEA analysis revealed that many datasets enriched in memory CD8^+^ T cells are enriched in WT early effectors (Fig. 2c, Table S2). Few of the analyzed datasets were enriched in *P2rx7*^-/-^ early effectors, with Bile Acid Metabolism being one of these (Fig. 2c, Table S2). Therefore, the presence of P2RX7 in early effectors is associated with simultaneous memory, stemness and residency transcriptional signatures.

**Figure 2.**
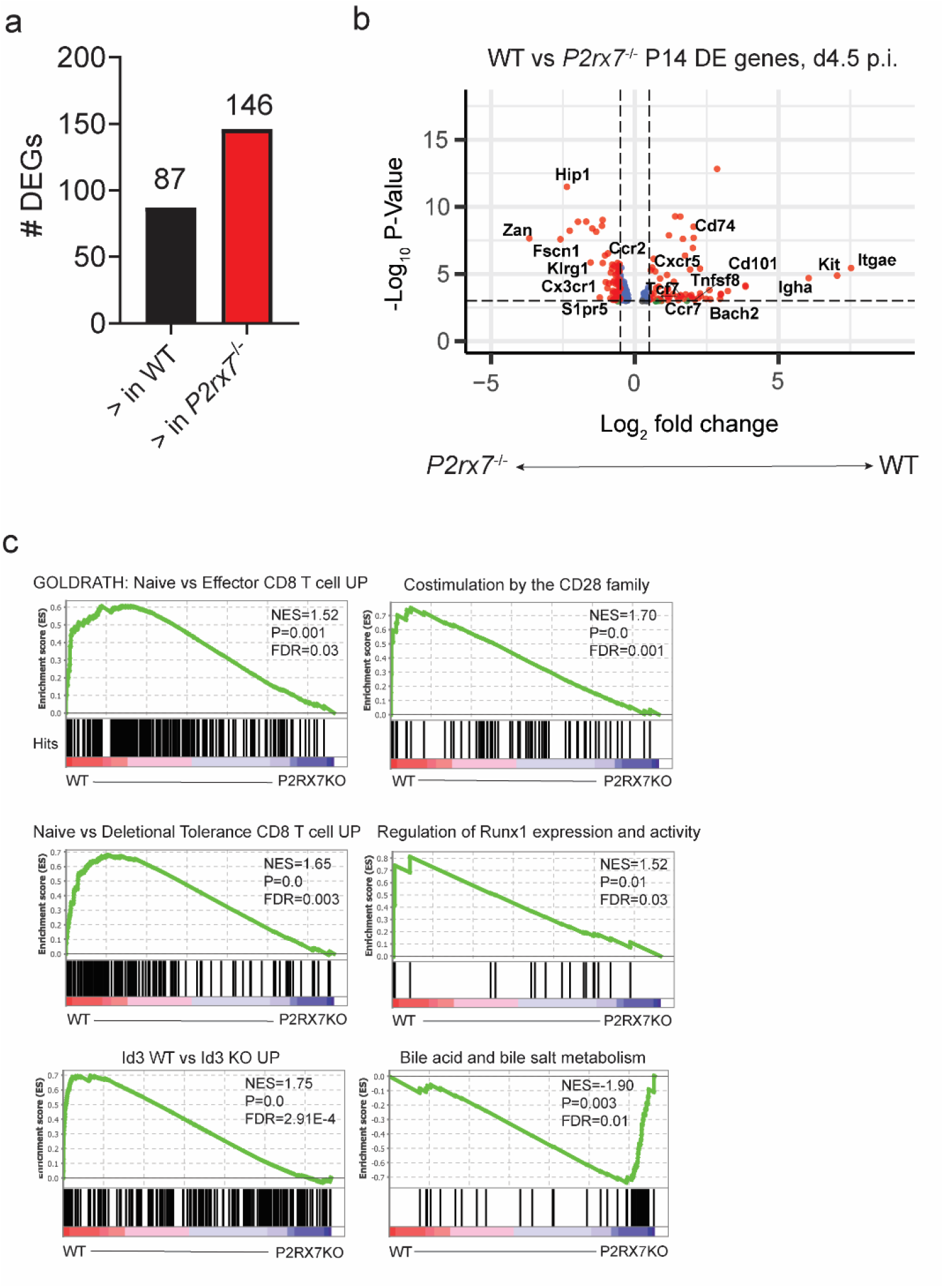
WT P14 cells are enriched for a pro-memory gene signature in comparison to *P2rx7*^-/-^ P14 cells. **(a-c)** Spleen WT and *P2rx7*^-/-^ P14 cells (previously co-adoptively transferred into recipient B6 mice) were sorted from recipient mice at 4.5 days after infection. The RNAs extracted from these populations were submitted for RNAseq analysis. (**a**) Bar graph showing the numbers of DEGs between WT and *P2rx7*^-/-^ P14 cells. (**b**) Volcano plot showing all DEGs upregulated (right) or downregulated (left) in WT P14 cells compared to *P2rx7*^-/-^ counterparts. Selected representative DEGs are denoted in this figure. (**c**) Gene Set Enrichment Analysis (GSEA) showing selected gene expression datasets enriched in WT or *P2rx7*^-/-^ (P2RX7KO) day 4.5 P14 cells. (**a-c**) Each replicate is a pool of spleen P14 cells from 4 mice, n=3 replicates per experimental group.

We next sought to define whether not only qualitative, but quantitative differences in P2RX7 expression would also differentially correlate with a pro-memory phenotype. We performed RNAseq analysis of P2RX7^hi^ versus P2RX7^lo^ early effector P14 cells, which cluster differentially (Fig. 3a) and yielded >400 DEGs (Fig. 3b-c, Table S3). Like the *WT*-*P2rx7*^-/-^ comparison, P2RX7^hi^ early effectors are enriched for memory CD8^+^ T cell genes, e.g. *Cxcr5, Id3, Xcl1, Ccr7, Tcf7* (Fig. 3c, Table S3), as well as Trm-specific genes like *Nr4a2* (Fig. 3c, Table S3), while P2RX7^lo^ early effectors are enriched either with TE genes (e.g. *Zeb2, Klrg1, Cx3cr1*) or anti-residency genes such as *Klf2* or *S1pr1* (Fig. 3b, Table S3). Again, this was reflected in the pathways enriched in P2RX7^hi^ or P2RX7^lo^ early effectors. While P2RX7^hi^ early effectors were enriched for many pathways associated with either survival or residency, P2RX7^lo^ early effectors were mainly enriched for apoptosis or terminal effector differentiation pathways (Fig. S2d, Table S2). This was also observed by GSEA analysis, where pro-memory datasets were enriched in P2RX7^hi^ early effectors and pro-TE phenotype or apoptosis-related datasets were enriched in P2RX7^lo^ cells (Fig. 3d, Fig. S2e, Table S2). Thus, not only P2RX7 presence but also its expression level is related to a simultaneous acquisition of a memory and residency transcriptional signatures in early effector CD8^+^ T cells. From these two analyses, we defined lists of genes that, at the early effector phase, are a) exclusively affected by P2RX7 presence, b) exclusively affected by P2RX7 expression level and c) affected by both P2RX7 presence and expression level (Fig. 3e, Table S4). Once again, among genes affected by both quantitative and qualitative P2RX7 differences, we found both memory versus TE-associated genes (e.g. *Ccr7, Tcf7, Zeb2, Cx3cr1*), and tissue-resident versus circulating CD8^+^ T cell-associated genes (e.g. *Nr4a2, Rgs10, Rgs16, S1pr1*) (Fig. 3e, Fig. S2f).

**Figure 3.**
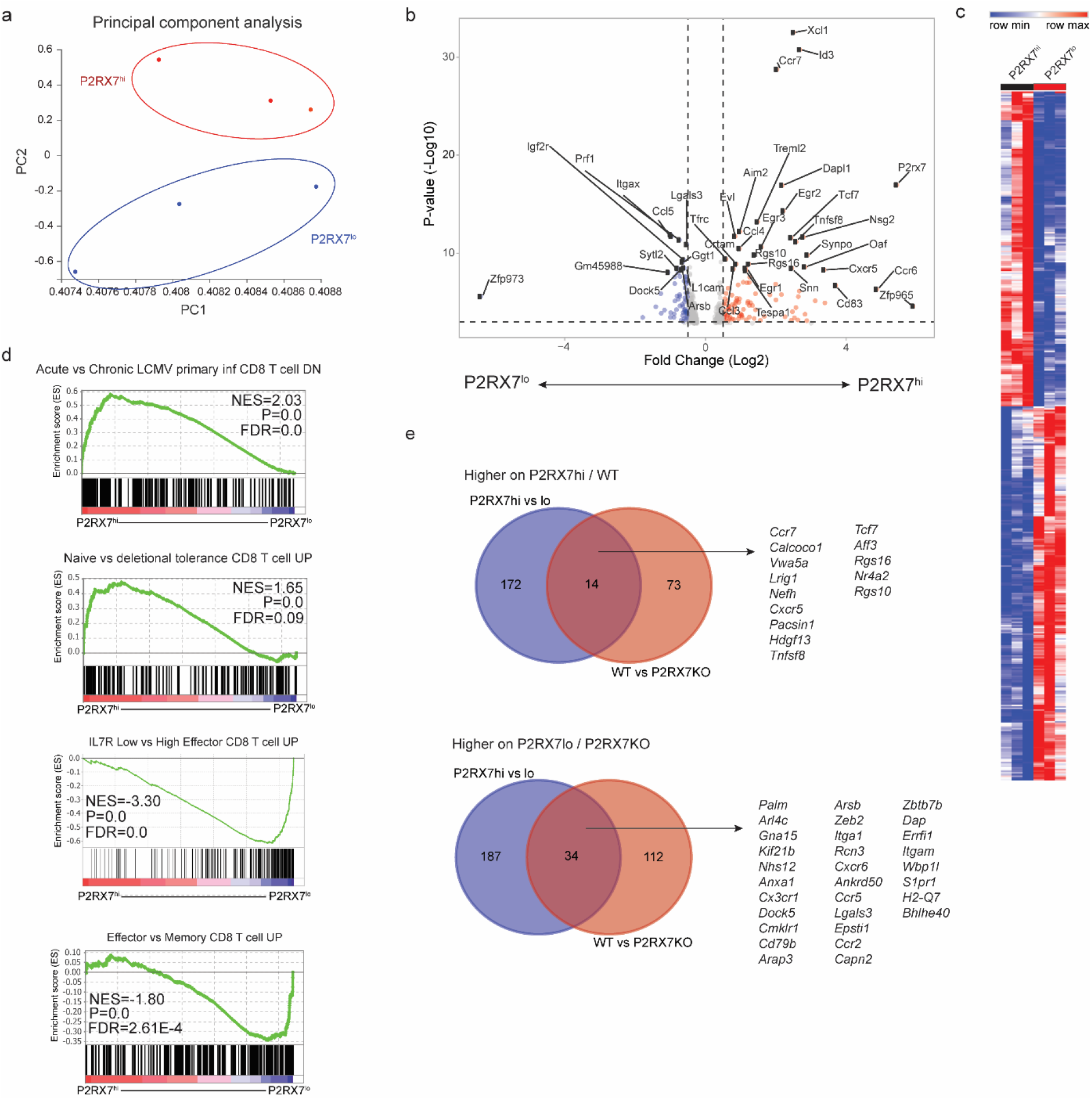
Defining the core genes quantitatively and qualitatively controlled by P2RX7 in early effector CD8^+^ T cells. **(a-d)** Spleen WT P14 cells (previously transferred into recipient B6 mice) were sorted from recipient mice at 4.5 days after infection based on 20% highest (P2RX7^hi^) and 20% lowest (P2RX7^lo^) expression. The RNAs extracted from these populations were submitted for RNAseq analysis. (**a**) Principal Component Analysis (PCA) plot showing the relation (based on PC1 and PC2 values) between P2RX7^hi^ and P2RX7^lo^ RNAseq samples. (**b**) Volcano plot showing all DEGs upregulated (right) or downregulated (left) in P2RX7^hi^ P14 cells compared to P2RX7^lo^ counterparts. Selected representative DEGs are denoted in this figure. (**c**) Heatmap showing the RNA count levels (normalized Z-scores) of all DEGs between P2RX7^hi^ and P2RX7^lo^ WT P14 cells. (**d**) Gene Set Enrichment Analysis (GSEA) showing selected gene expression datasets enriched in P2RX7^hi^ or P2RX7^lo^ day 4.5 P14 cells. (**e**) Venn Diagrams showing the correlation between DEGs found between WT and *P2rx7*^-/-^ day 4.5 P14 cells (red) and P2RX7^hi^ vs ^lo^ day 4.5 P14 cells (blue). The common DEGs found for both these comparisons (purple intersections) are listed. (**a-d**) Each replicate is a pool of spleen P14 cells from 6 mice, n=3 replicates per experimental group.

Next, we assessed the protein levels of many of the DEGs found in the RNAseq analyses. In early effector spleen P14 cells, a small proportion of cells express high levels of CD62L (Fig. S3a). A t-SNE analysis revealed that, among early effectors, highest expression of P2RX7 correlates with CD62L, and inversely correlates with KLRG1 (Fig. S3a). Indeed, P2RX7^hi^ early effector P14 cells display increased levels of the memory-associated molecules CD62L, CXCR5, and CD101 (and a trend in increased CD103), when compared to P2RX7^lo^ early effectors (Fig. S3b). Additionally, CD44, CD8α and the inhibitory molecule PD1 are also increased in P2RX7^hi^ early effectors (Fig. S3b). In counterpart, P2RX7^lo^ early effector P14 cells displayed enhanced expression of KLRG1 and Tim3 (Fig. S3b). Overall, we observed that P2RX7 heterogeneous expression in early effector CD8^+^ T cells associates with the simultaneous acquisition of a Tcm/Trm protein and transcriptional signatures.

### High P2RX7 expression favors the establishment of long-lived memory CD8^+^ T cells

Our transcriptional and protein data suggests that preferential P2RX7 expression, among early effectors, would favor the establishment of both Tcm and Trm CD8^+^ T cells. To investigate this hypothesis, we sorted P2RX7^hi^ and P2RX7^lo^ early effector spleen P14 cells, and adoptively transferred these cells into infection-matched secondary hosts (Fig. 4a). In synchrony with our previous data, P2RX7^hi^ progeny more efficiently established memory populations in the spleen (Fig. 4b), although a preferential enrichment of Tcm cells was not initially observed (Fig. 4b). The Tcm progeny from P2RX7^hi^ early effectors partly downregulated P2RX7 (Fig. 4c), although its average population expression was still substantially higher than the P2RX7^lo^ progeny, which never upregulated P2RX7 (Fig. 4c-d). At a late memory time point, we observed a further separation between the numbers of P2RX7^hi^ and P2RX7^lo^ progeny populations, and at this point a preferential enrichment of P2RX7^hi^-derived Tcm cells was observed (Fig. 4e). The P2RX7 bifurcation in early effectors also affected their ability to establish Trm populations, as observed in the SI IEL (Fig. 4f), Liver (Fig. S4a) and SG (data not shown). Similar to Tcm populations, we also observed an increased decay of P2RX7^lo^ progenies in the SI IEL Trm pool at late memory (Fig. 4f), although no differences in P2RX7 expression were observed between SI IEL transferred cells (Fig. S4b). The differential decay of P2RX7^hi^ versus P2RX7^lo^ early effector progenies are better exemplified in Fig. 4g. These results indicate that high P2RX7 expression in early effector CD8^+^ T cells preferentially favors the generation of long-lived Tcm and Trm cells.

**Figure 4.**
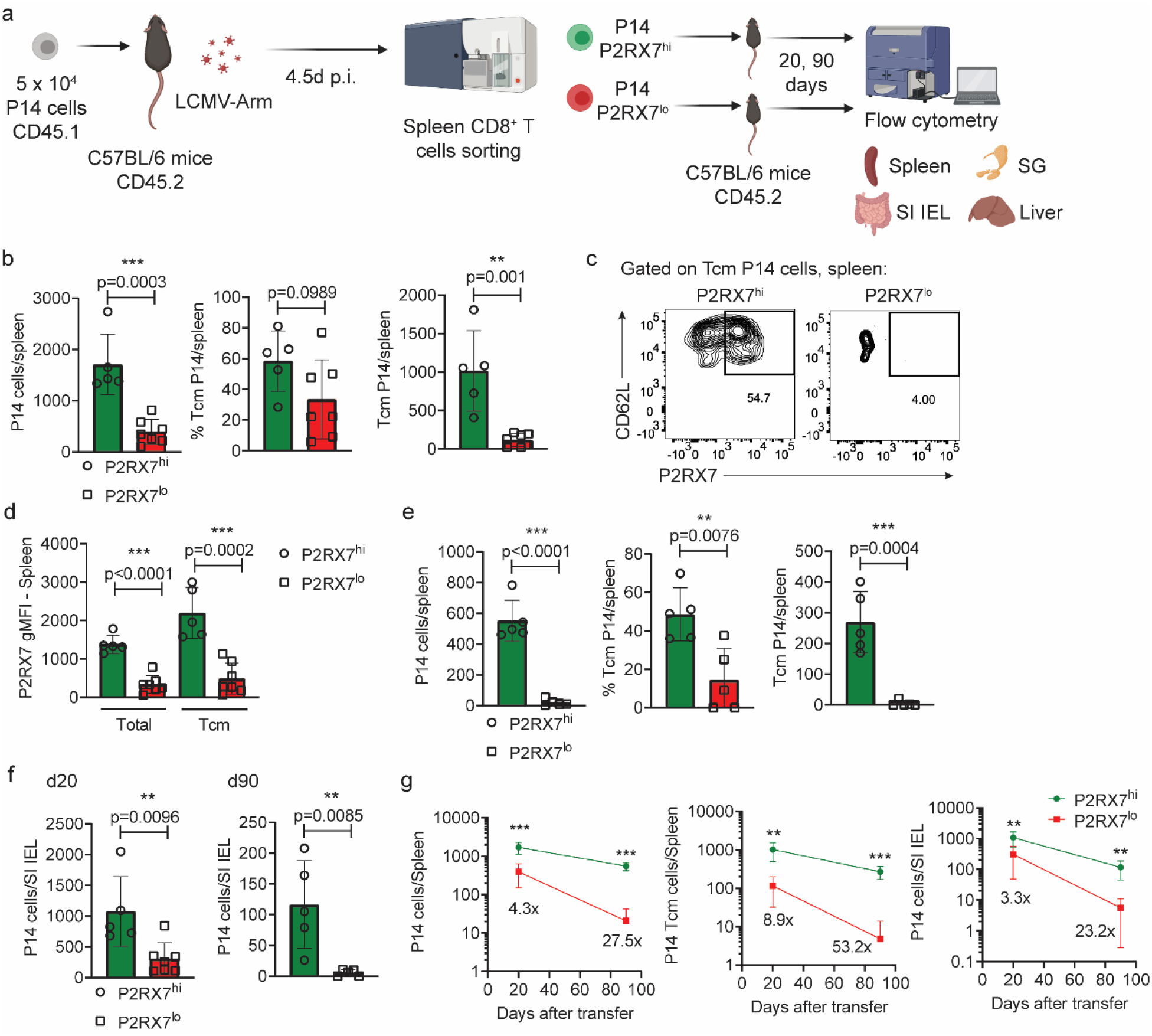
P2RX7 expression in early effectors favors the establishment of long-lived circulating and resident memory CD8^+^ T cells. (**a-g**) P14 cells were adoptively transferred into B6 mice (infected with LCMV-Arm), and at day 4.5 post-infection (p.i.), P2RX7^hi^ (20% highest) and P2RX7^lo^ (20% lowest) were sorted and adoptively transferred into infection-matched mice (3 × 10^5^ cells/mouse). At days 20 and 90 post-transfer, secondary recipient mice were assessed for transferred P14 cell numbers and phenotype. (**a**) Experimental plan for the secondary transfer experiments. (**b**) Total numbers (left), percentages of Tcm (center) and numbers of Tcm (right) P14 cells from P2RX7^hi^ or P2RX7^lo^ early effector progenitors at day 20 post-transfer. (**c**) Representative flow cytometry plots showing P2RX7 expression in Tcm P14 cells from P2RX7^hi^ and P2RX7^lo^ early effectors, at day 20 post-transfer. (**d**) Average P2RX7 gMFI levels in total and Tcm spleen P14 cells from P2RX7^hi^ and P2RX7^lo^ donors at day 20 post-transfer. (**e**) Total numbers (left), percentages of Tcm (center) and numbers of Tcm (right) P14 cells from P2RX7^hi^ or P2RX7^lo^ early effector progenitors at day 90 post-transfer. (**f**) Numbers of small intestine intraepithelial (SI IEL) P14 cells at day 20 (left) and day 90 (right) post-transfer. (**g**) Kinetics of the numbers of total spleen (left), spleen Tcm (center) and SI IEL (right) P14 cells over time after secondary transfer. The fold differences between P2RX7^hi^ and P2RX7^lo^ donor P14 cells are shown. (**b, d-g**) - Unpaired t-test, * - p<0.05, ** - p<0.01 and *** - p<0.001. (**b-g**) Average values ± SD, data pooled from 3 independent experiments.

### *Zeb2* knockdown preferentially rescue the memory CD8^+^ T cell pool in P2RX7-deficient CD8^+^ T cells

Among the genes affected by quantitative and qualitative P2RX7 expression differences in early effector CD8^+^ T cells, we found *Zeb2*, which encodes the zinc-finger transcriptional repressor Zeb2. This protein, which is expressed at higher levels in TEs relative to MPs (Table S1, Fig. S2f), promotes the TE phenotype at the expense of the memory CD8^+^ T cell pool^31,32, 33^. Given *Zeb2* was higher in P2RX7-deficient or P2RX7^lo^ early effector CD8^+^ T cells, we speculated that *Zeb2* ablation would preferentially rescue memory CD8^+^ T cell generation in the absence of P2RX7. To test this, we used RNP-based CRISPR-Cas9 to knockout *Zeb2* in WT or P2RX7-deficient P14 cells (Fig. 5a). *Zeb2* knockout, as previously described^31,32, 33^, significantly repressed the ability of WT P14 cells to differentiate into KLRG1^+^ early effectors or TEs at peak effector phase (Fig. 5b-c, 5e, Fig. S5a). *Zeb2* knockout in WT P14 cells promoted the MP phenotype and, after the early effector phase, the percentage of CD62L^+^ P14 cells (Fig. 5d-e, Fig. S5b), but did not lead to any numerical expansion of these populations (Fig. 5d, S5c). In contrast, *Zeb2*-ablated P2RX7-KO P14 cells not only showed a diversion toward the MP phenotype in terms of percentage, but also displayed a significant numerical increase in MPs (Fig. 5b-e, S5a-c). This scenario was also observed at the memory time point. In WT P14 cells, *Zeb2* knockout induced a mild increase in the percentage of Tcm populations (and a decrease in LLECs) (Fig. 5f, S5d), but this was not accompanied by changes in the number of memory P14 cells observed (Fig. 5f, S5c). *Zeb2*-ablated P2RX7-KO P14 cells, in contrast, not only underwent more profound alterations in the proportions of Tcm and LLECs observed (Fig. 5f, S5d), but also accumulated in higher numbers, such that there was no longer a statistical difference between WT and P2RX7-KO populations in the frequency of Tcm (Fig. 5f, S5c). *Zeb2* knockout, therefore, led to a disproportionate rescue of the ability of P2RX7-deficient CD8^+^ T cells to differentiate into circulating memory cells, especially the Tcm pool.

**Figure 5.**
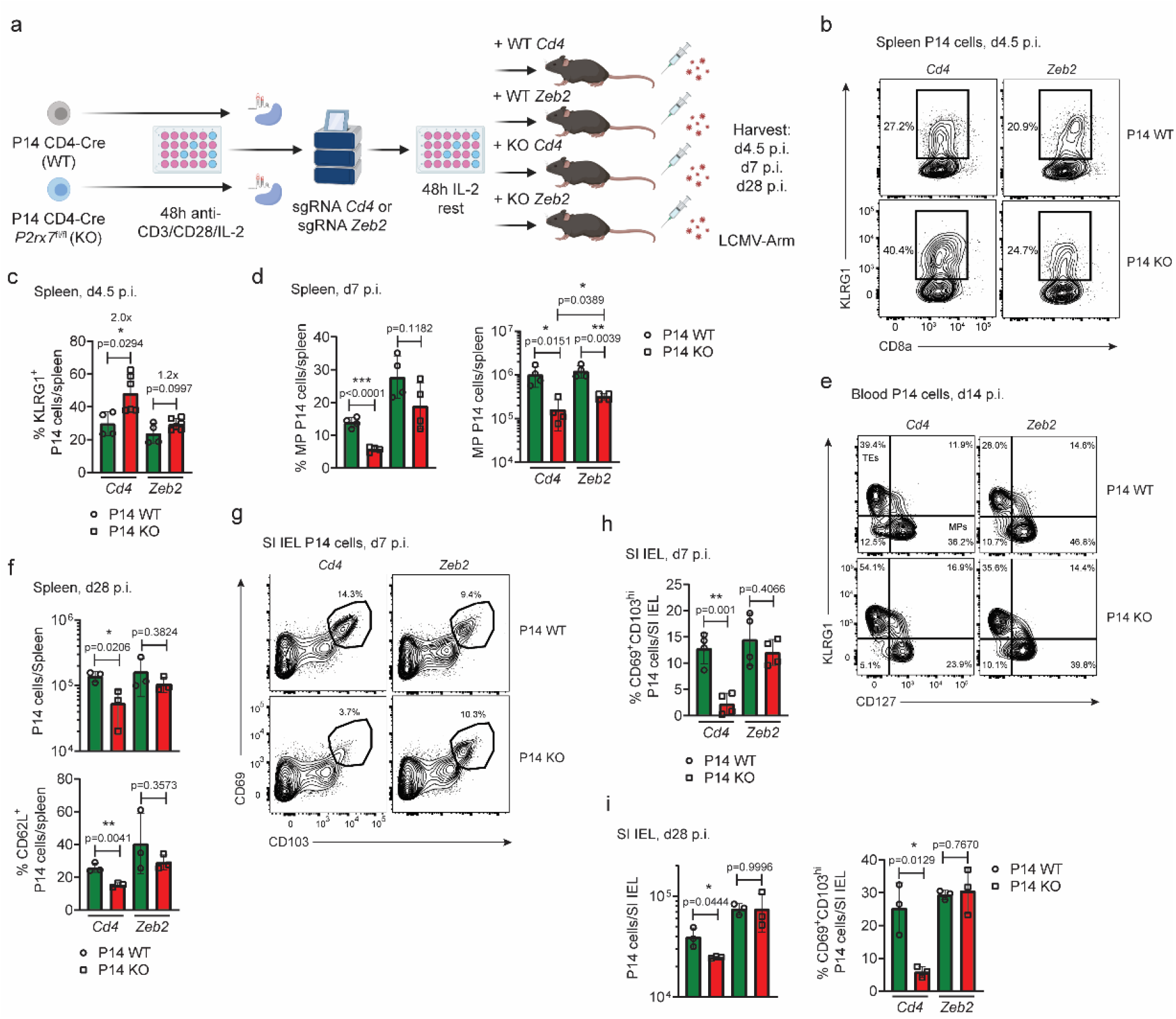
*Zeb2* knockout partially rescues the ability of P2RX7-deficient CD8^+^ T cells to form early memory precursors, Tcm and Trm subsets. (**a-i**) WT (CD4-Cre) and P2RX7-KO (CD4-Cre *P2rx7*^fl/fl^) P14 cells were in vitro activated; after 48h, CRISPR-Cas9 RNP-based knockdown of *Zeb2* or *Cd4* (control) were done. Cells were rested in IL-2 for additional 48h and subsequently transferred into recipient B6 mice. Recipient mice were infected with LCMV-Arm and the numbers and phenotype of transferred P14 cells were assessed over time after infection. (**a**) Experimental strategy of the CRISPR-Cas9 knockdown experiments. (**b**) Representative flow cytometry plots showing expression of KLRG1 in spleen P14 cells at day 4.5 after infection. (**c**) Percentages of spleen KLRG1^+^ P14 cells at day 4.5 after infection. (**d**) Percentages (top) and numbers (bottom) of KLRG1^-^CD127^+^ MP P14 cells in the spleen at day 7 after infection. (**e**) Representative flow cytometry plots showing expression of KLRG1 and CD127 in blood P14 cells at day 14 after infection. (**f**) Total numbers (top) and percentages of Tcm (CD62L^+^; bottom) spleen P14 cells at day 28 after infection. (**g**) Representative flow cytometry plots showing the expression of CD69 and CD103 in SI IEL P14 cells at day 7 after infection. (**h**) Percentages of CD69^+^CD103^hi^ SI IEL P14 cells at day 7 after infection. (**i**) Total numbers (top) and percentages of CD69^+^CD103^hi^ (bottom) SI IEL P14 cells at day 28 after infection. (**c-d, f, h-i**) – Unpaired t-test, * - p<0.05, ** - p<0.01 and *** - p<0.001. (**b-i**) Average values ± SD, data pooled from 2 independent experiments.

When Trm generation in the presence of *Zeb2* ablation was assessed, even more pronounced differences were observed. The initial tissue infiltration in SI IEL by total P14 cells was not affected by either P2RX7 knockout^24^ and/or *Zeb2* ablation (Fig. S5e). However, the differentiation of CD69^hi^CD103^hi^ cells, which we previously identified to be largely absent in P2RX7-KO cells^24^, was almost completely rescued when *Zeb2* was ablated (Fig. 5g-h). This was also observed at memory time points, where a numerical rescue of P2RX7-KO SI IEL Trm generation was also observed when *Zeb2* knockdown occurs (Fig. 5i). The infiltration and accumulation of P2RX7-KO SG P14 cells were also rescued by *Zeb2* knockdown (Fig. S5f-g). Overall, our data strongly indicates that P2RX7 promotes systemic virus-specific Tcm and Trm cells through inducing a pro-memory signature that is, at least partly, dependent on *Zeb2* negative regulation. It is of note that Zeb2, although a transcriptional repressor, did not directly affect P2RX7 expression in either circulating or Trm populations (Fig. S5h-j).

## Discussion

Sensing of microenvironmental changes plays key roles in the activation of many immune cells. CD8^+^ T cells are not different, as they express many receptors for extracellular signals. That includes, evidently, cytokine and chemokine receptors^18, 34^. Additionally, receptors for extracellular molecules released during infection are also expressed by CD8^+^ T cells during immune responses. One example is eATP, which can be released actively through exporting channels such as Pannexin-1^35, 36^, or passively by dying cells – due to local or systemic inflammation and tissue damage^22^. One eATP sensor in particular (P2RX7) is expressed at high levels and plays relevant roles in controlling CD8^+^ T cell responses^23, 24, 37, 38^. Because eATP can be released actively or passively, it is possible that CD8^+^ T cells are exposed to this signal not only in sites of active viral infection, but also in secondary lymphoid organs during initial effector function. Agreeing with this notion, P2RX7 is quickly upregulated in early effector CD8^+^ T cells^24^. In our study, we further explored this and found that P2RX7 upregulation in some early effectors favors the acquisition of transcriptional and protein memory signatures shared by Tcm and Trm cells. These findings suggest that the simultaneous promotion of Tcm and Trm cells by P2RX7 can be explained, at least in part, by differential sensing of eATP by early effector CD8^+^ T cells, which promote the generation of early memory precursors among some of these cells.

Previous reports have suggested that a minor fraction of early effector CD8^+^ T cells display heightened ability to differentiate into long-lived memory cells^12^, ^13^, ^14^. These studies mostly focused on the transcriptional regulation of this subset and have defined an important role for many transcriptional factors, especially TCF-1^13, 15^. These studies have also identified many receptors that, regardless of being controlled by TCF-1 or not, are associated with the memory fate. Whether these receptors for extracellular signals actively play a role in the early differentiation into memory precursors is unclear. Our results show that P2RX7 is one of these receptors, and actively promotes the diversion into the memory phenotype. Expression of P2RX7 and TCF-1 correlates in both CD8^+^ T cell early effectors and MPs. The hierarchy between TCF-1 and P2RX7 expression, however, is currently unclear. P2RX7 is upregulated in effector CD8^+^ T cells prior to TCF-1 re-expression^24^, which at first suggests P2RX7 antecedes TCF-1. Yet, a report using chronic LCMV suggests TCF-1 can bind to the *P2rx7* enhancer locus in CD8^+^ T cells^39^. Alternatively, a positive feedback mechanism may occur. In this situation, P2RX7 signaling in some early effectors may help the re-expression of TCF-1, and simultaneously TCF-1 reinforces the preferential upregulation of P2RX7. Future research will be needed to better define whether and how TCF-1 re-expression solidifies the preferential P2RX7 expression in early memory precursors.

Our data shows that differences in P2RX7 expression between CD8^+^ T cells occurs at the early effector phase, and it favors the later accumulation of memory CD8^+^ T cells. Our current data tracked these differences at the earliest time where peak P2RX7 expression in effector CD8^+^ T cells happens – at day 4-5 after LCMV infection^24^. However, P2RX7 upregulation can be detected as early as 2 days after LCMV or *in vitro* activation^24^. It is possible, therefore, that asymmetrical distribution of P2RX7 at even earlier time points may help direct the memory versus terminal effector programs in CD8^+^ T cells. Many studies have reported that asymmetrical partitioning of proteins in CD8^+^ T cells early after antigen priming favors the generation of memory cells in some daughters in detriment of the others^40, 41, 42, 43^. More specifically, asymmetric inheritance of TCF-1, T-bet and mTORC1 were all associated with these outcomes^16, 17, 41,44^. Aside the possible connection between TCF-1 and P2RX7, in our previous study we have shown P2RX7 expression in memory CD8^+^ T cells is inversely correlated with mTOR pathway activation^23^. Whether this is also true at the beginning of immune responses remains to be defined and may offer further mechanistic basis for the differentiation of P2RX7^hi^ CD8^+^ T cells in early memory precursors.

In contrast with the potential for early asymmetrical division to differentially promote memory versus terminal effectors, other reports suggest that memory CD8^+^ T cells arise from dedifferentiated effector CD8^+^ T cells^45, 46^. This apparent discrepancy may, on one side, reflect distinct experimental approaches or antigen stimulation nature. Alternatively, they may reflect heterogeneity in the memory CD8^+^ T cell populations formed in response to a determined antigen. Our results, indeed, showed that even P2RX7^lo^ early effectors or MPs can form memory populations, although less efficiently. Given P2RX7^lo^ early effectors display a pro-TE phenotype, these results suggest that even some TE-leaning CD8^+^ T cells can revert their phenotype towards memory. Therefore, our division between P2RX7^hi^ versus P2RX7^lo^ early effectors may define cells that develop into memory through early differentiation events and cells that develop into memory via dedifferentiation during T cell contraction.

Another unclear point is the dynamic changes we observed in P2RX7 expression after the early effector phase. The progeny from P2RX7^lo^ early effectors found in the circulation never upregulated P2RX7, even the few that developed into Tcm. In contrast, P2RX7^lo^ MPs were able to upregulate P2RX7. This suggests a complex expression control network, where low expression of P2RX7 at the early effector phase is accompanied by modifications that prevent its upregulation later. A possibility is that differential expression of P2RX7 is related to or promote epigenetic modifications in its own locus. Epigenetic modifications have been described to imprint numerous phenotypes in effector versus memory CD8^+^ T cells, with potential roles attributed to Tet and HDAC epigenetic modulators^47, 48, 49^. Of note, HDAC-related gene sets were enriched in P2RX7^lo^ early effector CD8^+^ T cells. Whether epigenetic alterations at the *P2rx7* locus explain this expression dynamics it remains to be defined. Nevertheless, this should not be related to P2RX7 expression in Trm cells, as even the few P2RX7^lo^ early effectors differentiating into Trm cells express high levels of P2RX7. In this case, other signaling pathways and transcriptional controllers may function in the *P2rx7* locus after non-lymphoid tissue entry, for example retinoic acid signaling^50^.

The acquisition of a Trm cell potential, although possibly requiring additional signals after non-lymphoid tissue entry^24,25,51^, is potentially imprinted in certain circulating early effector CD8^+^T cells^52^. The nature of this imprint is not clear, however. It may, on one side, rely on stochastic signals sensed by a subset of early effectors regardless of TCR differences^11^. Alternatively, it may reflect distinct TCR functional avidities and/or affinities as recently suggested^52^. The extracellular signals promoting this imprinting are also not fully explored. Our previous report suggested that P2RX7-mediated eATP sensing may favor the subsequent generation of Trm precursors^24^. Here, we not only confirmed these findings, but also provided additional evidence that P2RX7 asymmetrical expression in early effectors already aligned with transcriptional differentiation involving certain residency-versus-circulation signatures. They do not exclude, by any means, the likelihood that eATP sensing through P2RX7 also plays a role in Trm generation after non-lymphoid tissue entry, which our previous report also suggested as a mechanism for the function of this receptor^24^. Rather, the presence of Trm cells from P2RX7-imprinted versus non-imprinted early effectors may reflect heterogeneous populations forming in non-lymphoid tissues. Our results showed that P2RX7^hi^-derived Trm cells tended to survive longer, which suggests P2RX7 signaling at the early effector phase promotes a longer-lived subset of Trm cells. In agreement with this notion, two recent reports have shown *P2rx7* differentially expressed in a SI IEL-infiltrating CD8^+^ T cell subset that also contained long-lived memory-associated genes^10, 51^. Whether P2RX7 signaling in early effectors preferentially gives rise to a long-lived subset of Trm cells remains to be defined and is a current goal of our group.

We have found that P2RX7 high expression at the early effector phase was negatively correlated with *Zeb2* expression. Zeb2 directly promotes the TE/LLEC phenotype at the expense of memory CD8^+^ T cell subsets, but without major effects in the total numbers of memory CD8^+^ T cells formed^31,33^. In contrast with these previous reports, *Zeb2* ablation not only re-directed the P2RX7-deficient CD8^+^ T cell pool toward memory differentiation and away from TEs, but also significantly increased the size of the memory cell population. These results suggest that Zeb2 expression is negatively controlled by P2RX7 upregulation. Moreover, they indicate that either Zeb2 function is somehow curbed by P2RX7 signaling, or that P2RX7 upregulation promotes indirect control of the Zeb2 expression and function. A possible candidate for this is TCF-1, which is positively correlated with P2RX7, negatively correlates with Zeb2 and is not directly controlled by Zeb2^31^. Two other signals that reportedly repress Zeb2 are TGF-β and mir-200 microRNAs^32^. Although we did not focus here on mir-200 factors, we did find an enrichment for the TGF-β signaling pathway in not only WT-versus-knockout early effectors – also previously reported by us^24^ – but also between P2RX7^hi^-versus-P2RX7^lo^ early effectors, suggesting an additional pathway potentially involved in the P2RX7-mediated control of Zeb2 function. The connection between P2RX7, TGF-β signaling and Zeb2 may also help explain why *Zeb2* ablation also led to significant numerical rescue of P2RX7-deficient Trm cell generation – although the enrichment in early memory precursors may also explain this result. In summary, this work highlights how differential acquisition of eATP sensing ability serves to transcriptionally promote many aspects of a pro-memory signature in a subset of early effector CD8^+^ T cells. Identifying the factors precisely controlled by P2RX7-eATP sensing in early effectors will allow significant improvement of immunizations against infection or cancer, including the use of T cell-targeted P2RX7 agonism as a form of promoting memory CD8^+^ T cells in the circulation and in barrier tissues simultaneously.

## Methods

### Mice

Female C57BL/6 (B6) and B6.SJL (expressing the CD45.1 allele) mice were purchased from the National Cancer Institute (via Charles River). *P2rx7*^-/-^ and CD4-Cre mice were obtained from Jackson Laboratories and were both fully backcrossed onto C57BL/6 background. P2rx7fl/fl mice were obtained from Drs. Gyorgy Hasko (Rutgers University) and Matyas Sandor (U-Wisconsin), crossed onto the C57BL/6 background and with CD4-Cre mice. LCMV-D^b^GP33-specific TCR transgenic P14 mice were fully backcrossed to B6, *P2rx7*^-/-^, CD4-Cre and CD4-Cre *P2rx7*^fl/fl^ mice, with introduction of CD45.1 and CD45.2 congenic markers for identification.

Animals were maintained under specific-pathogen-free conditions at the University of Minnesota and in Mayo Clinic Arizona. In all experiments, mice were randomly assigned to experimental groups. All experimental procedures were approved by the institutional animal care and use committee at the University of Minnesota (IACUC 1709-35136A) or at Mayo Clinic Arizona (IACUC A00005542-20).

### Viral strains

LCMV Armstrong strain was maintained at −80°C until infection and diluted to 2 × 10^6^ PFU/ml in PBS.

### Infection studies

Wild-type, *P2rx7*^-/-^ or CD4-Cre *P2rx7*^fl/fl^ P14 cells were adoptively transferred into naive wild-type mice, which were infected with LCMV Armstrong strain (2 × 10^5^ PFU, intraperitoneally (i.p.)).

### Primary cell cultures

Naive P14 cells were obtained from six- to 8-week old male or female mice with a C57BL/6 background. P14 cells were cultured in complete RPMI media: RPMI 1640 (Corning) supplemented with 10% Fetal Bovine Serum (Atlanta Biologicals), 100 U/ml penicillin/streptomycin (Thermo Fisher Scientific) and 2 mM L-glutamine (Corning). All cells were cultured at 37°C in a humidified atmosphere containing 5% CO_2_.

### Adoptive cell transfer

CD8^+^ P14 cells were negatively enriched (using the CD8+ T cell isolation kit – Miltenyi Biotec) from WT (CD45.1/2^+^) and *P2rx7*^-/-^ (CD45.2/2^+^) P14 mice. The cell populations were mixed at a ratio of 1:1 and a total of 5 x 10^4^ P14 cells were co-adoptively transferred into recipient mice (CD45.1/1^+^), which were infected the next day with LCMV-Arm. At appropriated times after infection, mice were sacrificed for analysis. In other experiments, WT (CD45.2/2^+^) or CD4-Cre *P2rx7*^fl/fl^ (CD45.2/2^+^) P14 cells were negatively enriched and individually adoptively transferred (5 x 10^4^ cells) into recipient mice (CD45.1/1^+^), which were infected with LCMV-Arm. For the secondary adoptive transfer experiments, CD45.2/2^+^ P2RX7^hi^ or P2RX7^lo^ MPs or day 4.5 effector P14 cells were sorted from the spleens of recipient mice, then adoptively transferred (3 × 10^5^ cells) into infection-matched recipient mice (CD45.1/1^+^). At the appropriate time points after transfer, mice were sacrificed for analysis.

### Flow cytometry

Lymphocytes were isolated from tissues including spleen, blood, small intestine epithelium (SI IEL), liver and salivary glands (SG) as previously described^30, 53^. In summary, organs were removed and cut in small pieces into erlenmeyers containing 30 mL of 0.5 mg/ml Collagenase type I solution (SG) or 0.15 mg/ml Dithioerythritol (SI IEL). Following this period, lymphocytes were isolated by 44/67% Percoll gradient isolation. During isolation of lymphocytes from non-lymphoid tissues, in all experiments, 50 μg of Treg-Protector (anti-ARTC2.2) nanobodies (BioLegend) were injected i.v. 30 minutes prior to mouse sacrifice^54^. Direct *ex vivo* staining and intracellular staining were performed as described^30, 55^. To identify LCMV-specific CD8^+^ T cells, PE- or APC-gp^33–41^ tetramers were obtained from the NIH tetramer core. For detection of vascular-associated lymphocytes in non-lymphoid organs, *in vivo* i.v. injection of PerCP-Cy5.5-conjugated CD8α antibody was performed^56^. Due to unclear division between bona-fide Trm and circulating memory using i.v. CD8α injection in the liver, we used total liver P14 numbers in all experiments where this organ is listed. Among LCMV-specific CD8^+^ T cells, the following markers were used to distinguish these respective populations: Tcm (CD44^+^CD62L^+^), Tem (CD44^+^CD62L^−^CD127^+^), LLEC (CD44^+^CD62L^−^CD127^−^KLRG1^hi^), Trm (i.v.CD8α^−^ CD69^+/−^CD103^hi/int/lo^), MPs (CD127^+^KLRG1^−^), TEs (CD127^−^KLRG1^+^). For detection of the intracellular factor TCF-1, surface-stained cells were permeabilized, fixed and stained by using the eBioscience Foxp3 staining kit, according to manufacturer instructions. Flow cytometric analysis was performed on LSR II or LSR Fortessa (BD Biosciences) and data was analyzed using FlowJo software (Treestar). A complete list of antibodies used for flow cytometry analysis is found in Table S4.

### Cell sorting

Cell sorting was performed on a FACS Aria III device (BD Biosciences). RNA expression experiments were performed with CD45.1/2^+^ Wild-type (CD44^+^) and CD45.2/2^+^ *P2rx7*^-/-^ (CD44^+^) P14 CD8^+^ T cells sorted from mice 4.5 days post-LCMV infection, or with CD45.2/2^+^P2RX7^hi^CD44^+^ and CD45.2/2^+^ P2RX7^lo^CD44^+^ P14 CD8^+^ T cells from mice 4.5 days post-LCMV infection. Secondary adoptive transfer experiments were done with P2RX7^hi^ and P2RX7^lo^KLRG1^−^CD127^+^ (MPs) P14 CD8^+^ T cells from mice 8 days post-LCMV infection, with CD62L^+^CD44^+^ (Tcm) P14 CD8^+^ T cells from mice 30 days post-LCMV infection, or with P2RX7^hi^CD44^+^ and P2RX7^lo^CD44^+^ P14 CD8^+^ T cells from mice 4.5 days post-LCMV infection. The population purity after cell sorting was > 95% in all experiments.

### RNA-seq and bioinformatics analysis

WT or *P2rx7*^-/-^ P14 cells sorted from spleen at day 4.5 post-LCMV infection were first homogenized using QIAshredder columns (Qiagen) and RNA was extracted using RNeasy Plus mini kit (Qiagen) following manufacturer’s instructions. Library preparation and RNA-seq libraries were prepared and RNA-seq (150-bp pair-end, NovaSeq 6000 Illumina) was done at the University of Minnesota Genomics Center. RNA-seq reads were mapped and counted by CHURP to generate the raw count matrix. The raw count values were converted to FPKM (Fragments Per Kilobase Million) values and were further transformed to z-score values to perform PCA analysis. Differential expressed gene (DEG) analysis was done using DESeq2^57^. Genes with more than 2-fold changes and FDR lesser than 0.05 were kept for gene cluster analysis.

P2RX7^hi^ or P2RX7^lo^ P14 cells sorted from spleen at day 4.5 post-LCMV infection were homogenized using QIAshredder and RNA extracted using RNeasy Plus mini kit as described before. Library preparation and RNA-seq (DNBseq platform, PE 100bp pair-end read length) was done by BGI Americas. RNA-seq reads were mapped and raw count matrix was generated. DEG analysis was done using DESeq2, and genes with >2-fold changes and FDR <0.05 were considered for gene cluster analysis.

Heatmaps were generated using the Morpheus software (Broad Institute - https://software.broadinstitute.org/morpheus/). R studio was used to generate Volcano plots of DEGs. The GSEA software was used to generate comparisons with established gene sets. For the P2RX7^hi^ versus P2RX7^lo^ analysis, BGI provided customized analysis of DEGs, PCA plots and GSEA analyses between groups.

### CRISPR-Cas9 experiments

Cas9/RNP nucleofection of P14 cells was performed as described previously^58^. Briefly, P14 cells were isolated and activated as described above. While cells were maintained in culture, single guide RNAs for *Zeb2* or *Cd4*, and Cas9 protein were mixed by pipetting up and down and pre-complexed at room temperature for at least 10 min. After this period, 1-10 million activated P14 cells were re-suspended in 20 μL primary cell nucleofection solution (Lonza), then mixed and incubated with the crRNA/Cas9 mix for 2 min at room temperature. The P14 cell/Cas9/RNP mixes were transferred to Nucleofection cuvette strips (4D-Nucleofector X kit S; Lonza). Cells were electroporated using a 4D nucleofector (4D-Nucleofector Core Unit; Lonza). After nucleofection, prewarmed complete RPMI was used to transfer transfected P14 cells in 96-well plates. After 2 h, P14 cells were cultured in 24-well plates in complete RPMI (+ IL-2 – 10 ng/ml) for 48 h, before adoptive transfer into recipient mice (1 × 10^5^ cells/mouse).

### Quantification and Statistical Analysis

Details on statistics used can be found in figure legends. Data were subjected to the Kolmogorov-Smirnov test to assess normality of samples. Statistical differences were calculated by using unpaired two-tailed Student’s t test (or one-way ANOVA with Tukey post-test, where indicated). All experiments were analyzed using Prism 7 (GraphPad Software). Graphical data was shown as mean values with error bars indicating the SD. P values of < 0.05 (*), < 0.01 (**) or < 0.001 (***) indicated significant differences between groups.

## Acknowledgments

We thank the members of the Borges da Silva and Jamequist lab for intellectual input. This work was funded by NIAID grants awarded to H.BdS. (R00 AI139381) and to S.C.J. (R01 AI038903, AI145147).

## Author contributions

H.BdS. conceived the project and designed the experiments. H.BdS. and S.C.J. provided mice and reagents for all experiments. T.V-K., S.vD., C.P., K.M.W. and H.BdS. performed the experiments. T.V-K., S.vD. and H.BdS. analyzed the data. H.BdS. wrote the manuscript, with input from all authors.

