## Supplementary Figures for "The extracellular ATP receptor P2RX7 imprints a pro-memory transcriptional signature in effector CD8^+^ T cells"

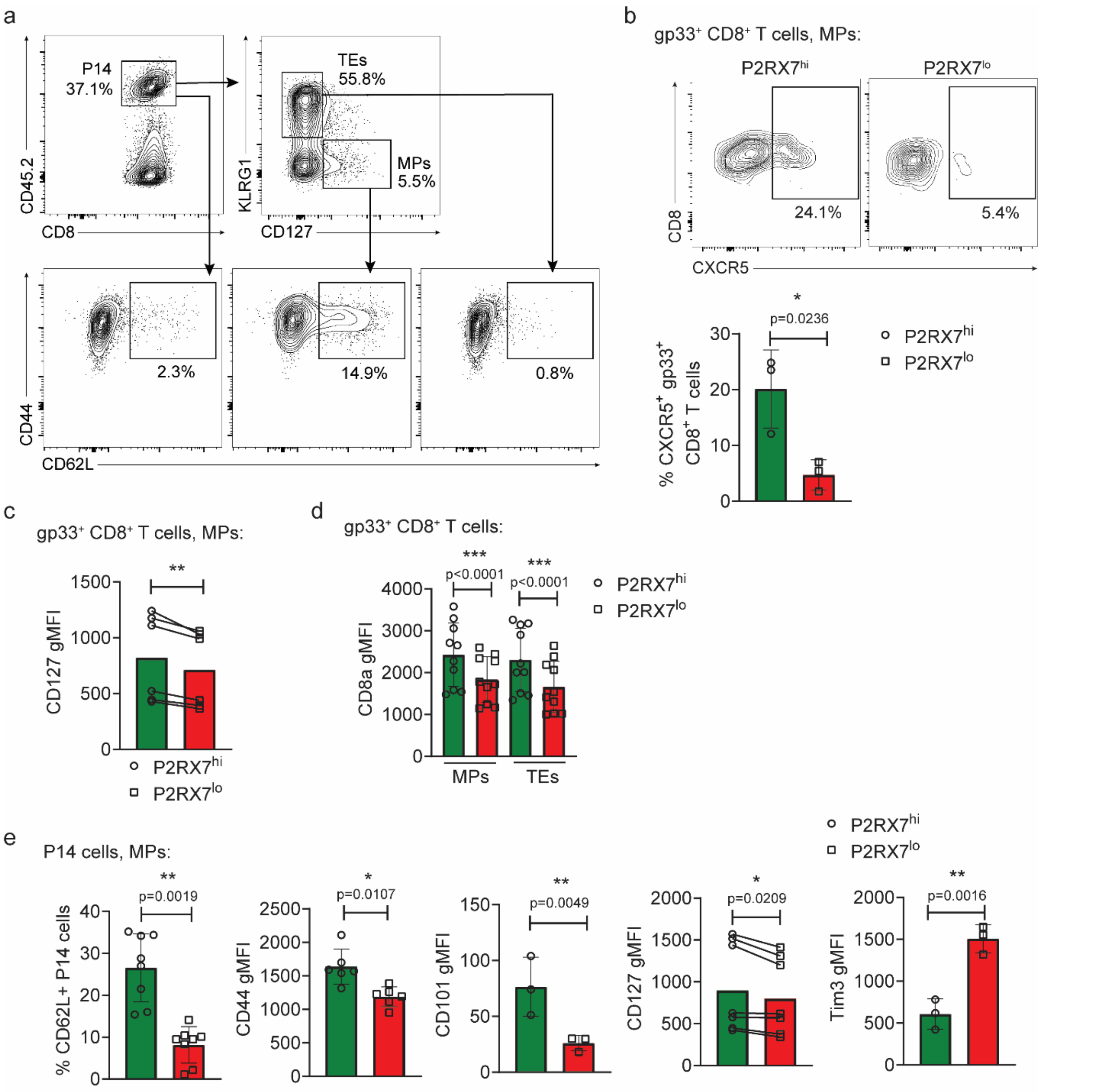


**Figure S1. Heterogeneity in memory markers between P2RX7$^{hi}$ vs P2RX7$^{lo}$ MP cells.** (a) Representative flow cytometry plots showing (in P14 cells) the gating strategy for the determination of TEs, MPs and Tcm precursors (CD62L$^{+}$ MPs). **(b-d)** C57BL/6 mice were infected with LCMV-Arm and harvested at 7 days post-infection; the protein expression of multiple molecules in antigen-specific (gp33$^{+}$) CD8$^{+}$ MPs (day 7 after infection) is shown. **(b)** Top: Representative flow cytometry plot showing expression of CXCR5 in gp33$^{+}$ P2RX7$^{hi}$ and P2RX7$^{lo}$ MPs. Bottom: Percentages of CXCR5$^{+}$ gp33$^{+}$ P2RX7$^{hi}$ and P2RX7$^{lo}$ MPs. (**c**) Average CD127 gMFI values in P2RX7$^{hi}$ and P2RX7$^{lo}$ gp33$^{+}$ MPs. (**d**) Average CD8α gMFI values in P2RX7$^{hi}$ and P2RX7$^{lo}$ gp33$^{+}$ MPs and terminal effectors (TEs). (**e**) In other experiments, P14 cells were adoptively transferred into B6 mice which were infected with LCMV-Arm. Phenotypic analysis of spleen P14 MPs were done at day 7 after infection. Average values for percentages of CD62L$^{+}$ P14 MPs and CD44/CD101/CD127/Tim3 gMFI in P14 MPs are shown. (**b-e**) – Unpaired t-test, * - p<0.05, ** - p<0.01 and *** - p<0.001. (**a-d**) Average values ± SD, data pooled from 3 independent experiments.

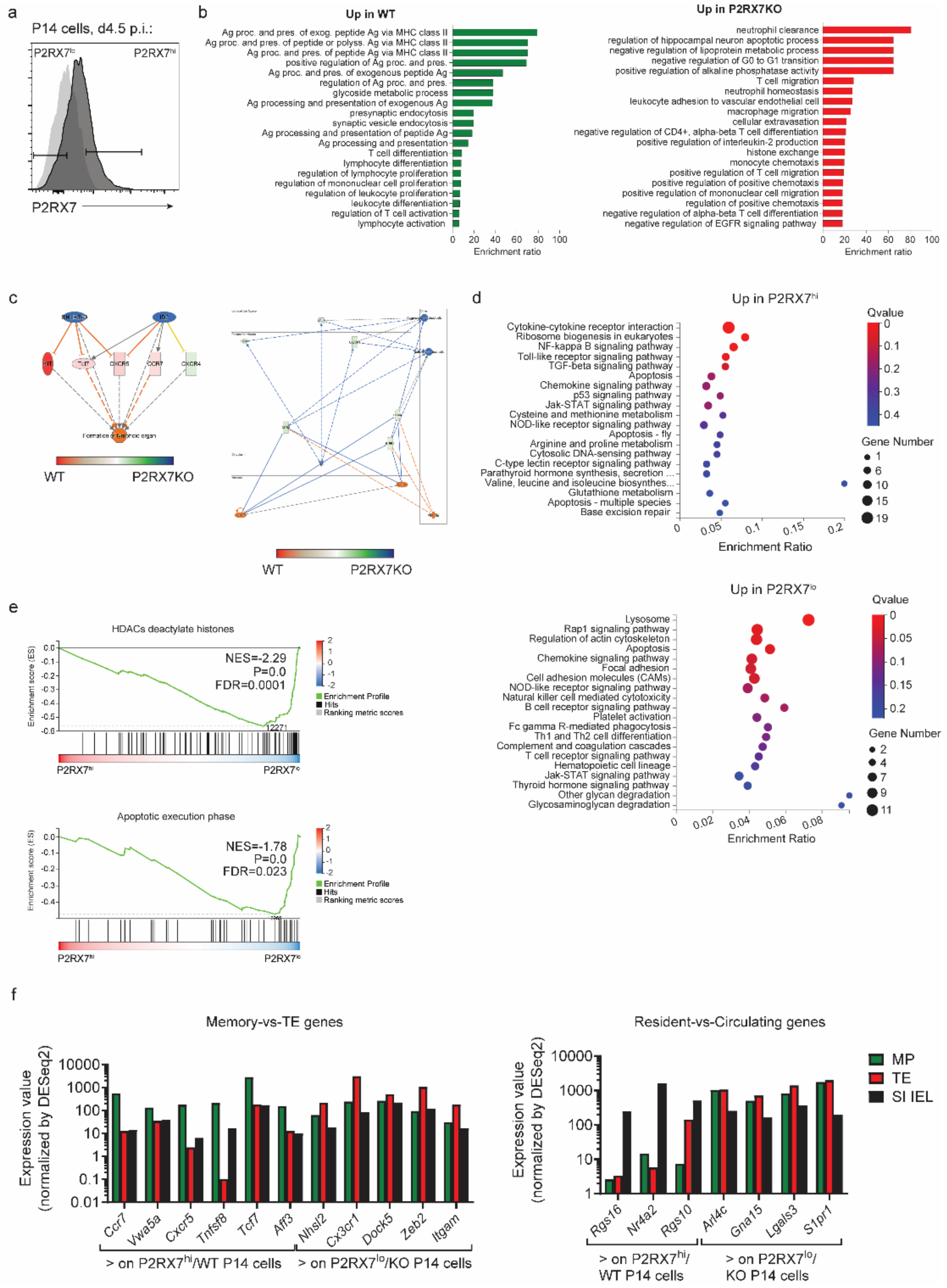


**Figure S2. Transcriptional signatures controlled by P2RX7 qualitatively and quantitatively in early effector CD8+ T cells.** (**a**) Representative flow cytometry histogram showing the expression of P2RX7 in P14 cells at day 4.5 post-LCMV (dark) and naïve (light grey). (**b-e**)

RNAseq analyses were done to assess the transcriptional signatures associated with WT vs *P2rx7*$^{-/-}$ (**b-c**) and P2RX7$^{hi}$ vs P2RX7$^{lo}$ (**d-e**) spleen P14 cells at day 4.5 post-LCMV infection. (**b**) Main gene ontology (GO) pathways enriched in WT (green) or *P2rx7*$^{-/-}$ (red) day 4.5 spleen P14 cells. Data shows enrichment ratio values per GO pathway. (**c**) Two examples of gene networks differentially affected in WT vs *P2rx7*$^{-/-}$ P14 cells; genes and/or networks enriched in WT P14 cells are depicted in red/orange/pink, and the ones enriched in *P2rx7*$^{-/-}$ P14 cells are depicted in blue/green. (**d**) Main GO pathways enriched in P2RX7$^{hi}$ (top) and P2RX7$^{lo}$ (bottom) day 4.5 P14 cells. Data shows enrichment ratio values, False Discovery rates (FDR) Q-values and number of genes representing each GO pathway. (**e**) Two representative GSEA analyses showing dataset gene enrichment in P2RX7$^{lo}$ day 4.5 P14 cells. (**f**) Immgen (Immgen.org) RNA expression values (day 7 MPs, TEs or SI-IEL P14) of selected genes enriched (in our analysis) in WT/P2RX7$^{hi}$ day 4.5 P14 cells or *P2rx7*$^{-/-}$/P2RX7$^{lo}$ day 4.5 P14 cells. (**b-c**) Each analysis replicate is pooled from 5 different recipient mice, n=3 replicates per experimental group. (**d-e**) Each analysis replicate is pooled from 6 different recipient mice, n=3 replicates per experimental group.

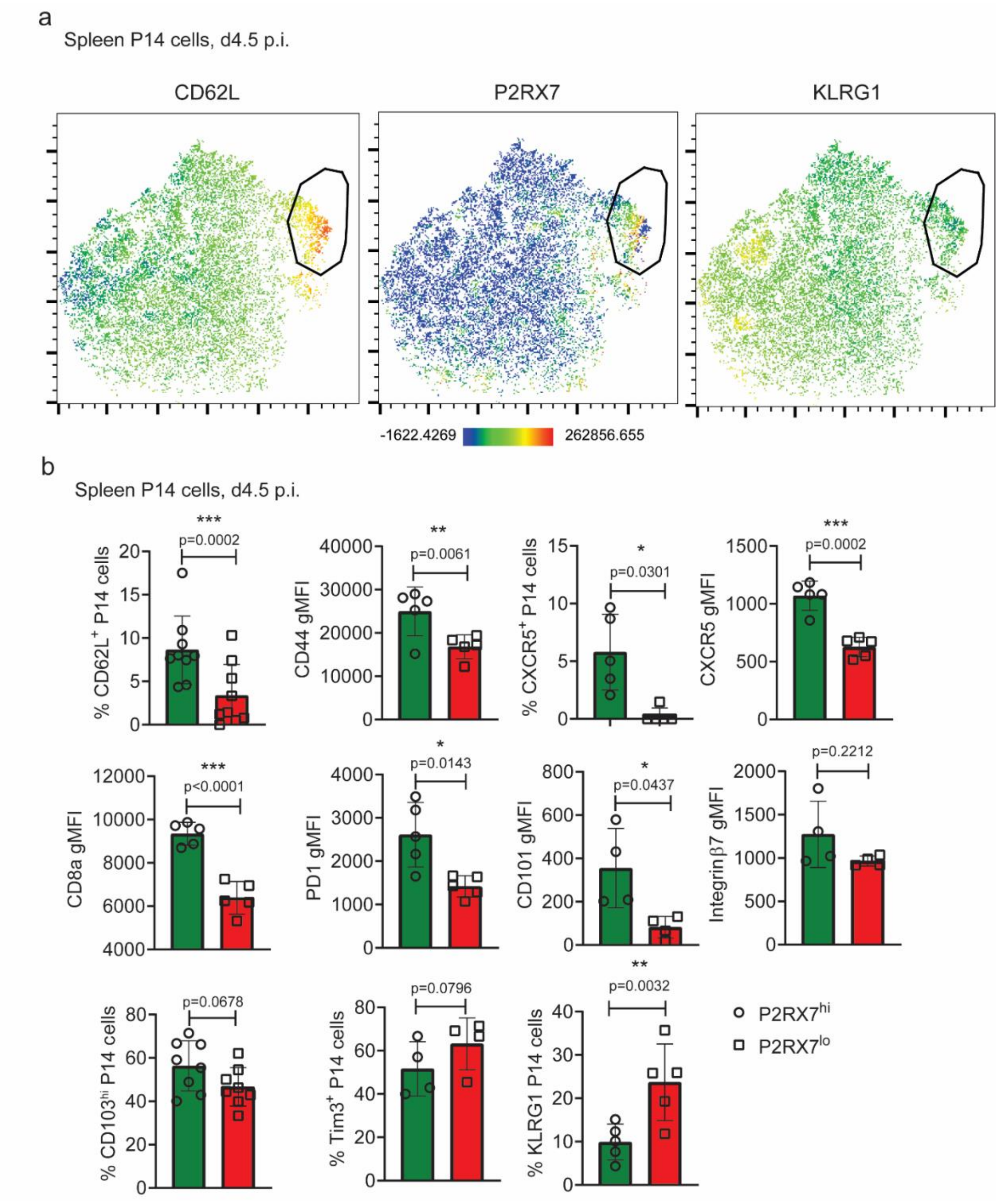


**Fig. S3. P2RX7 protein expression in early effector CD8$^+$ T cells correlates with memory markers expression.** (**a-b**) WT P14 cells were adoptively transferred into recipient B6 mice, which were infected with LCMV-Arm. Infected mice spleens were harvested at day 4.5 post-infection for P14 cell analysis. (**a**) Representative t-SNE plots depicting the protein expression of CD62L, P2RX7 and KLRG1 among spleen P14 cells. (**b**) Average values for percentages of CD62L$^+$/CXCR5$^+$/CD103$^{hi}$/Tim3$^+$/KLRG1$^+$, and CD44/CXCR5/CD8α/PD1/CD101/Integrin β7 gMFI in P2RX7$^{hi}$ and P2RX7$^{lo}$ spleen P14 cells. (**b**) – Unpaired t-test, * - p<0.05, ** - p<0.01 and *** - p<0.001. (**b**) Average values ± SD, data pooled from 2-3 independent experiments.

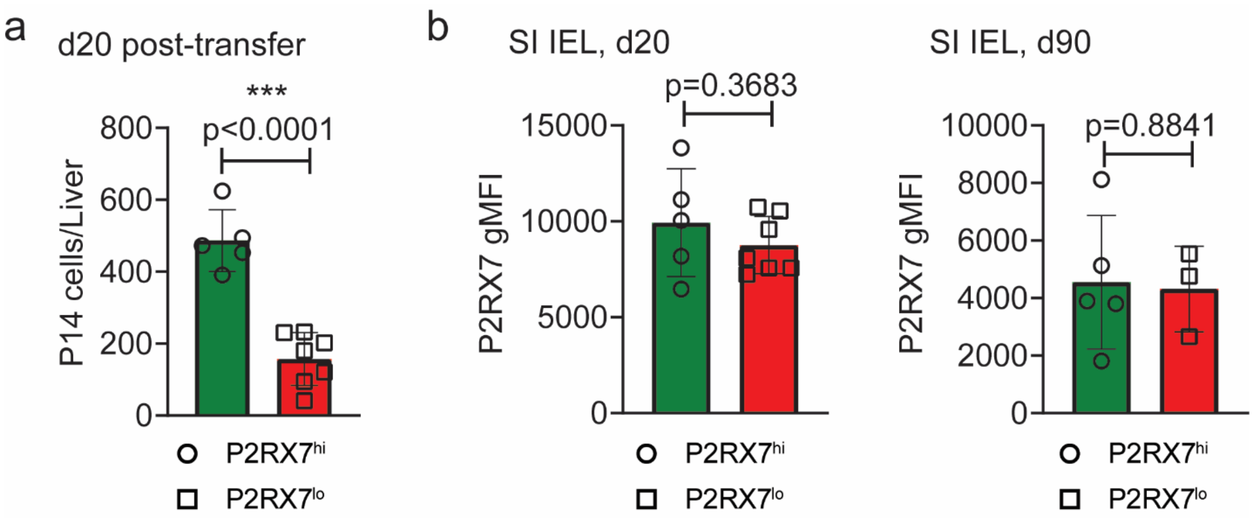


**Fig. S4. P2RX7$^{hi}$ early effectors preferentially form Trm cells but P2RX7 expression in Trm cells is upregulated in both P2RX7$^{hi}$ and P2RX7$^{lo}$ progenies.** (**a-b**) P14 cells were adoptively transferred into B6 mice (infected with LCMV-Arm), and at day 4.5 post-infection, P2RX7$^{hi}$ (20% highest) and P2RX7$^{lo}$ (20% lowest) P14 cells were sorted and adoptively transferred into infection-matched mice (3 x $10^5$ cells/mouse). At days 20 and 90 post-transfer, secondary recipient mice were assessed for transferred P14 cell numbers and phenotype. (**a**) Average numbers of P14 cells in the liver at 20 days post-transfer. (**b**) Average P2RX7 gMFI values in SI IEL P14 cells at 20- and 90-days post-transfer. (**a-b**) – Unpaired t-test, * - p<0.05, ** - p<0.01 and *** - p<0.001. (**a-b**) Average values ± SD, data pooled from 2-3 independent experiments.

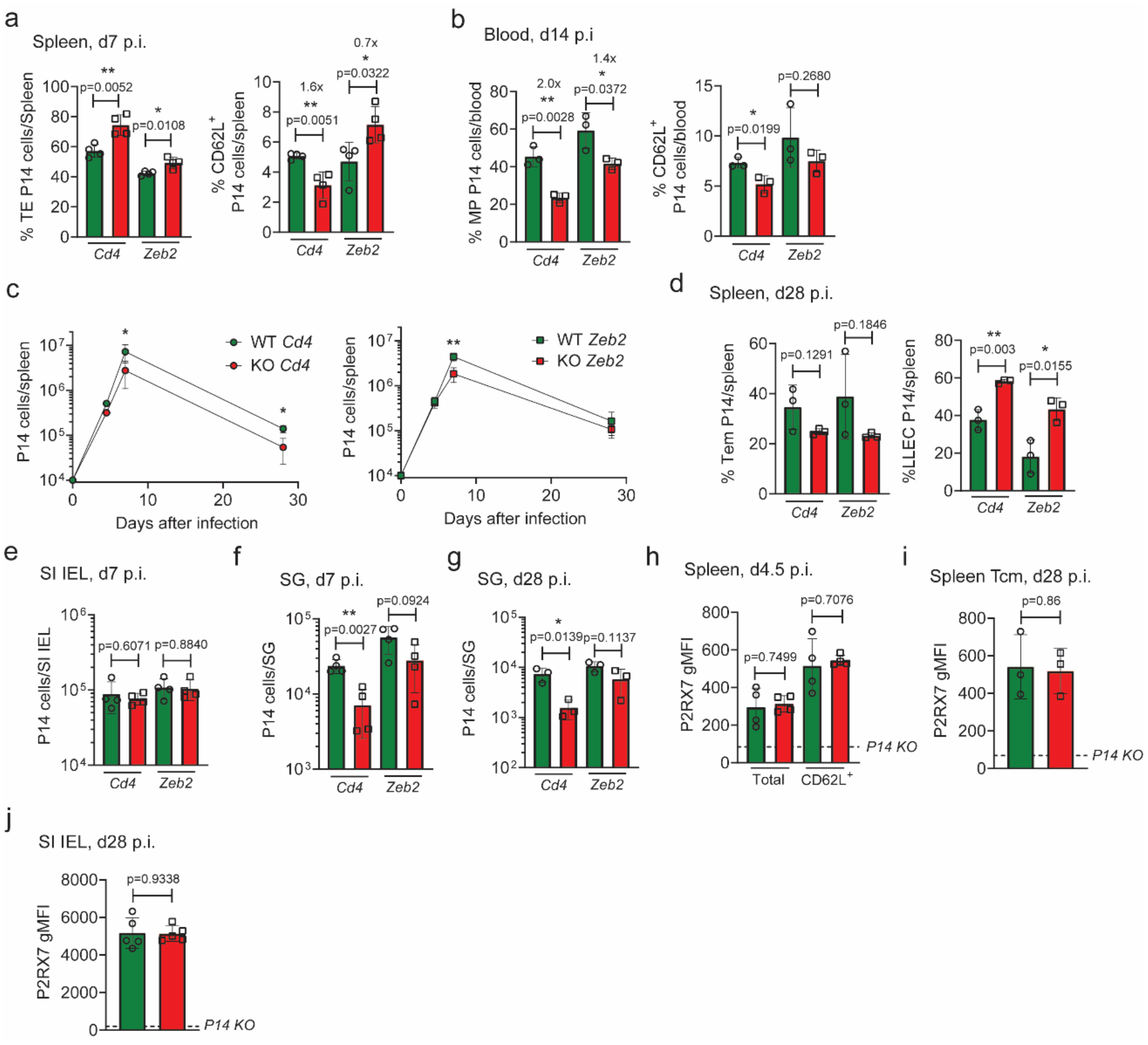


**Fig. S5. *Zeb2* knockdown significantly rescues the ability of P2RX7-deficient CD8⁺ T cells to differentiate into Tcm and Trm populations.** (**a-j**) WT (CD4-Cre) and P2RX7-KO (CD4-Cre *P2rx7*$^{fl/fl}$) P14 cells were *in vitro* activated; after 48h, CRISPR-Cas9 RNP-based knockdown of *Zeb2* or *Cd4* (control) were done. Cells were rested in IL-2 for additional 48h and subsequently transferred into recipient B6 mice. Recipient mice were infected with LCMV-Arm and the numbers and phenotype of transferred P14 cells were assessed over time after infection. (**a**) Percentages of terminal effectors (TEs) and CD62L$^+$ spleen P14 cells at day 7 after infection. (**b**) Percentages of MP (left) and CD62L$^+$ (right) blood P14 cells at day 14 after infection. (**c**) Numbers of spleen WT/P2RX7-KO P14 cells over time after infection with *Cd4* sgRNAs (left) and *Zeb2* sgRNAs (right). (**d**) Percentages of Tem (left) and LLEC (right) spleen P14 cells at day 28 after infection. (**e**) Numbers of SI IEL P14 cells at day 7 after infection. (**f**) Numbers of salivary gland (SG) P14 cells at day 7 after infection. (**g**) Numbers of SG P14 cells at day 28 after infection. (**h**) Average P2RX7 gMFI values in total and CD62L$^+$ spleen P14 cells at day 4.5 after infection. (**i**) Average P2RX7 gMFI values in spleen Tcm P14 cells at day 28 after infection. (**j**) Average P2RX7 gMFI values in SI IEL P14 cells at day 28 after infection. (**a-j**) – Unpaired t-test, * - $p<0.05$, ** - $p<0.01$ and *** - $p<0.001$. (**a-j**) Average values ± SD, data pooled from 2-3 independent experiments.
